# Transcriptome Landscape of Human Oocytes and Granulosa Cells Throughout Folliculogenesis

**DOI:** 10.1101/285445

**Authors:** Yaoyao Zhang, Zhiqiang Yan, Qingyuan Qin, Vicki Nisenblat, Yang Yu, Tianren Wang, Cuiling Lu, Ming Yang, Shuo Yang, Ying Yao, Xiaohui Zhu, Xi Xia, Yujiao Dang, Yixin Ren, Peng Yuan, Rong Li, Ping Liu, Hongyan Guo, Jinsong Han, Haojie He, Yu Wu, Meng Li, Kun Zhang, Yiting Wang, Jie Qiao, Jie Yan, Liying Yan

**Affiliations:** Center for Reproductive Medicine, Department of Obstetrics and Gynecology, Peking University Third Hospital, No.49 North HuaYuan Road, HaiDian District, Beijing 100191, China; National Clinical Center for Obstetrics and Gynecology, Beijing 100191, China; Key Laboratory of Assisted Reproduction, Ministry of Education, Beijing 100191, China; Beijing Key Laboratory of Reproductive Endocrinology and Assisted Reproduction, Beijing 100191, China; Peking-Tsinghua Center for Life Sciences, Peking University, Beijing 100871, China; Department of Obstetrics and Gynecology, Peking University Third Hospital, No.49 North HuaYuan Road, HaiDian District, Beijing 100191, China; Beijing Advanced Innovation Center for Genomics (ICG), Peking University, Beijing 100871, China

## Abstract

Folliculogenesis is a highly regulated process that involves bidirectional interactions of the oocytes and surrounding granulosa cells (GCs). Little is unknown, however, about the transcriptomic profiles of human oocytes and GCs throughout folliculogenesis. Here we performed a high resolution RNA-Seq of human oocytes and GCs at each follicular stage, which revealed unique transcriptional profiles, stage-specific signature genes, oocyte- and GC-derived genes that reflect ovarian reserve. We identified reciprocal cell-to-cell interactions between oocytes and GCs, including NOTCH, TGF-β signaling and gap junctions and determined the expression patterns of maternal-effect genes involved in folliculogenesis and early embryogenesis. Finally, we demonstrated robust differences between human and mice oocyte transcriptomes. This is the first comprehensive overview of the transcriptomic signatures governing the stepwise human folliculogenesis in-vivo that provides a valuable resource for basic and translational research in human reproductive biology.

## INTRODUCTION

Human folliculogenesis is a remarkably complex, well-orchestrated process that relies on a synchronization between oocyte maturation and proliferation of the neighboring granulosa cells (GCs) ^1, 2^. Follicle growth and oocyte maturation are associated with dynamic transcriptional events in both oocyte and GC compartments of the follicle, featured by high transcriptional activity of growing follicles post recruitment and transcriptional silencing of mature oocytes ^3^. Despite the impressive body of data produced in recent years on oocyte biology, many questions regarding the key differentiation events occurring during folliculogenesis in humans remain unanswered Oocyte-GCs bidirectional communications via signal transduction or direct cell-to-cell contact provide molecular and structural basis for effective oocyte-GC crosstalk required for adequate follicular development. It is poorly understood, however, what initiates ^4^ the communication between human follicle compartments and how this interaction is regulated and maintained.

Genes that accumulate in oocytes during maturation and influence early embryonic development are referred to as maternal-effect genes^5^. The events associated with maternal-effect genes and the process of maternal-to-zygotic transition have been studied in animal models ^6^, albeit the expression pattern and the role of these genes during oocyte development in humans remain unclear. In women, reproductive potential and reproductive lifespan depend on the follicle number available for fertilization and the rate of follicular loss during reproductive years, both of which determine ovarian reserve ^7^. Even though the levels of tests such asanti-Mullerian hormone and follicle stimulating hormone (FSH), and the antral follicle count measured by transvaginal ultrasound has been widely used to predict ovarian reserve, the universally accepted markers that accurately predict fertility potential and ovarian reserve are still lacking ^8–10^.

Till now, most data on follicular transcriptome are derived from the studies on animal models due to restricted availability of human follicles for research. While animal studies have advanced our understanding of ovarian biology, expression patterns of oocyte-specific genes are known to display considerable interspecies variations ^11^. In humans, transcriptomic studies of ovarian follicle rely on evaluations of the oocytes at limited stages of development or on analysis of unseparated follicle compartments ^12–14^, which undermines the applicability of the findings to understanding of a coherent whole of folliculogenesis. Application of single-cell RNA sequencing (scRNA-Seq) and other high resolution sequencing techniques allows an unbiased evaluation of cells of any size and efficient amplification of low-abundant transcripts, which makes these methods a powerful tool in studying the biological process of interest^15^. Therefore, characterizing human oocyte and GC transcriptome during different stages of development at high resolution is key to understanding the events that coordinate human oocyte maturation and follicle growth, which in turn is expected to provide remarkable opportunities for developing novel diagnostic and therapeutic approaches to improve fertility.

This work aimed at analyzing the gene expression patterns throughout folliculogenesis by exploring the transcriptome of human oocytes and GCs at five key stages of follicular development in-vivo, from primordial to preovulatory stage. Using scRNA-Seq approach, we identified dynamic expression patterns with distinct transcriptome signatures in oocytes and GCs, recognized the oocyte-GC communications during folliculogenesis and revealed putative biomarkers to predict ovarian reserve. In addition, we investigated the expression patterns of maternal-effect genes in human oocytes and propose their role in follicular development. Finally, we compared our data with the published oocyte transcriptome profiles from rodents and demonstrated robust interspecies differences.

## RESULTS

### Global Transcriptome Profiling of Human Oocytes and GCs

A total of 83 oocytes and 92 GC samples (each GC sample comprised randomly selected 10 cells per follicle owing to low abundancy of RNA in these cells) were obtained (Figure 1A). The data for MII oocytes were downloaded from our previous research^13^ and merged with the newly generated data for the subsequent analyses. A down sampling analysis revealed adequate sequencing depth for gene expression detection (Figure S1D). After quality control and filtration of the RNA-Seq data, 80 oocytes and 71 GC samples were retained for the analysis (Figures S1). The unsupervised principal component analysis (PCA) showed there was a clear separation between 80 oocytes and 71 GCs with distinction between the developmental stages (Figure 1B). Next we analyzed the expression patterns of the known cell-type markers in the oocyte and GC clusters (Figure 1C and Figure 1D), which confirms the validity of PCA classification. Moreover, we selected the most variable genes that contributed to PC1 and could distinguish between the oocytes and GCs. In oocytes, the most variable genes included *NOL4*, *CTCTL*, *ITIH2*, *SNAP91* and *NAALAD2*. In GCs, the most variable genes were: *ZEB2*, *CD44*, *HSPG2*, *KDSR* and *SRRM3*The top oocyte or GCs-expressed variable genes ate presented in Figures 1E and 1F, respectively. the above observations suggest that the most variable genes between oocytes and GCs could be candidate cell type-specific markers.

**Figure 1.**
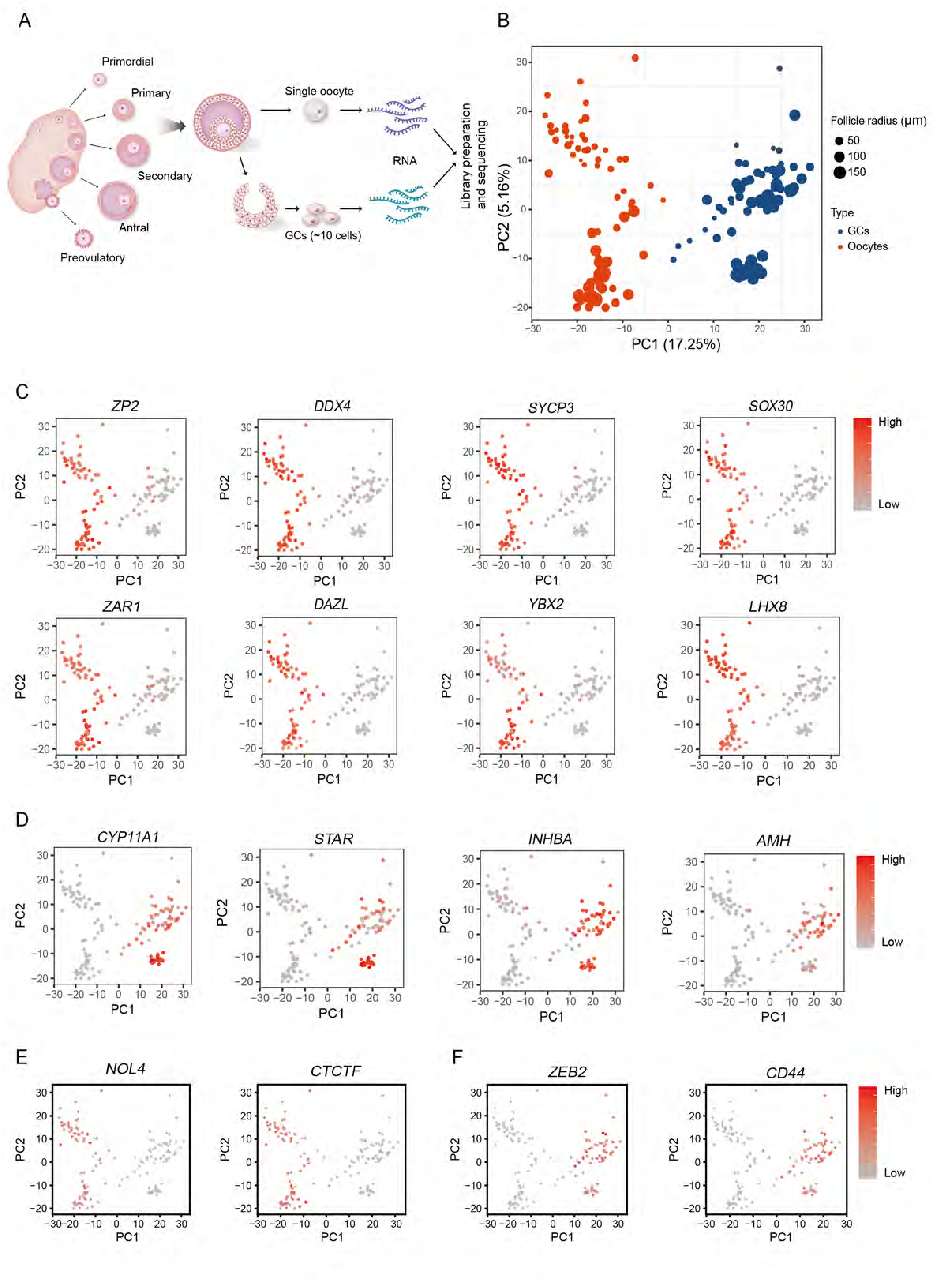
The Global Transcriptome Patterns of Human Oocytes and granulosa cells (GCs). A) Schematic illustration of the study workflow. Primordial - primordial follicles; Primary - primary follicles; Secondary - secondary follicles; Antral - antral follicles; Preovulatory - preovulatory follicles; GCs - granulosa cells. B) Principal component analysis (PCA) of the transcriptome of scRNA-Seq data from all oocytes and GCs included in this study. The PC1 separates two follicular compartments (oocytes vs. GCs). The PC2 separates the samples according to their follicular stages. Red color indicates oocytes, blue color indicates GCs. The sizes of the points represent follicles of different sizes. C) Expression patterns of oocyte marker genes exhibited on PCA plots; a gradient of gray to red indicates the low to high gene expression level. D) Expression patterns of GC marker genes exhibited on PCA plots. E) Expression of the candidate cell-type specific markers of oocytes on PCA plots. F) Expression of the candidate cell-type specific markers of GCs on PCA plots.

### Gene Expression Dynamics and Transcriptional Characteristics of Oocytes during Folliculogenesis

To explore the gene expression patterns in oocytes during folliculogenesis, we performed PCA analysis within the cohort of oocytes and observed five distinct subpopulations which corresponded morphological classification of follicular development (Figure 2A). The stage of follicular development was much more powerful discriminator than the sampling entity (Figure S1E), indicating that transcriptome profiles of oocytes reflect the physiological status of maturation, rather than genetic background. We then characterized the differentially expressed genes (DEGs) between the stages of follicle development in oocytes (FDR < 0.05; FC of log_2_ transformed FPKM > 1.5) (Figure 2B). A large number of these genes were under-expressed in primordial follicles with progressively increasing number of activated transcripts with follicular growth and maturation. The expression profiles of DEGs *RBM24*, *GPD1, NTF4* and *LCP2* were validated by using immunohistochemistry (IHC) on ovarian tissue from the scRNA-Seq sample set. Results showed that *RBM24* was expressed only in primordial and primary stage, *GPD1* was restricted secondary and antral stage, while *NTF4* and *LCP2* were specifically expressed in antral stage. These results are in agreement with the scRNA-Seq results (Figure 2E). To investigate possible biological functions of DEGs involved in folliculogenesis, we performed Gene Ontology (GO) analysis of DEGs in oocytes and revealed enriched biological processes in association with each stage of follicular development (*p* < 0.05) (Figure 2C).

**Figure 2.**
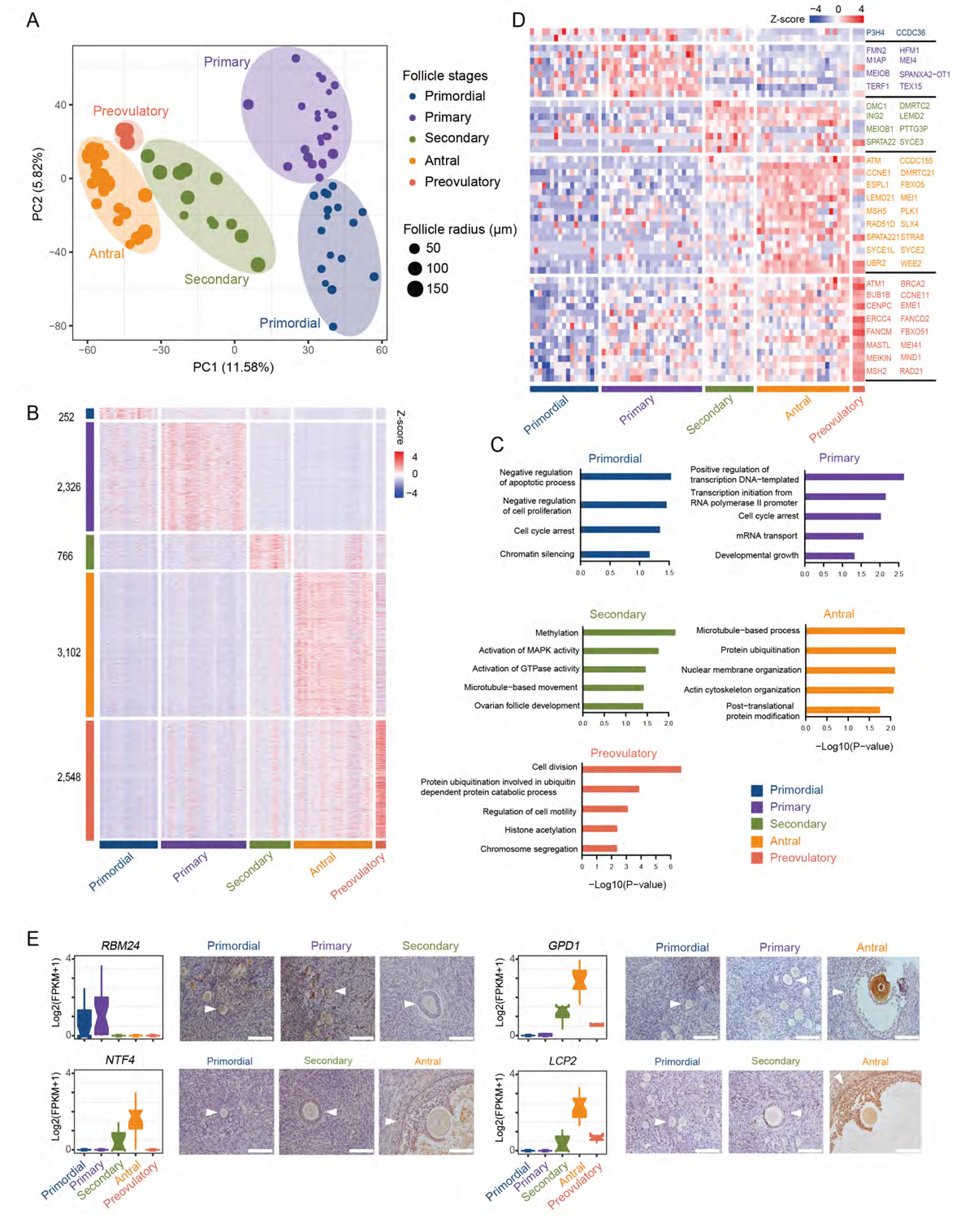
Gene Expression Dynamics and Transcriptional Characteristics of Oocytes. A) Principal component analysis (PCA) of the transcriptome of scRNA-Seq data from 80 oocytes collected from follicles at different stages of development. Oocytes are clustered into five subpopulations corresponding to morphological stages. Different colors and sizes of the points represent follicles of different stages and sizes, respectively. See also Figure S1E for the results of PCA that portrays sampling entities within the oocytes. B) Heatmap of all the differentially expressed genes (DEGs) in oocytes at five stages of folliculogenesis. The numbers of identified DEGs are indicated on the y-axis, and the stages of follicle development are presented along the x-axis. The color key from blue to red indicates the relative gene expression level from low to high, respectively. C) Significantly enriched GO terms (biological processes) of DEGs in oocytes at five stages of folliculogenesis. D) Heatmap of genes involved in meiosis that are differentially expressed in oocytes at five stages of folliculogenesis. E) Immunohistochemistry staining of selected DEGs. The boxplot demonstrating gene expression level is presented on the left of each corresponding immunohistochemistry panel. White triangles indicate the follicles. Scale bar, 100 µm.

We also investigated a number of germ cell markers and revealed some exhibited stable expression levels in oocytes during follicle development (e.g. *DDX4* and *ZP3*, Figure S2A), while others showed progressive upregulation with oocyte maturation (e.g. *ZP1, GDF9* and *H1FOO*). The expression of *ZP2*, *ZP3* and *ZP4* remained stable across all stages. Our findings suggest that *ZP1* might play a role in acquisition of oocyte competence and *ZP2*, *ZP3* and *ZP4* may act across all developmental stages in human oocytes. In contrast with our previous findings in fetal ovary ^16, 17^, this study showed that the expression levels of pluripotency markers *NANOG* and *POU5F1* were constantly low during oocyte development, suggesting their different roles in fetal and adult ovary.

Next, we focused on investigating the expression patterns of meiosis-related genes in oocytes. The sets of 195 genes involved in meiosis I and 13 genes involved in meiosis II were gathered from the GO Consortium. Of these, 52 genes were differentially expressed across different stages of oocyte maturation. Most of the meiosis-specific genes demonstrated shift towards upregulation as maturation of oocytes proceeded, and exhibited strong over-expression in antral and preovulatory follicles (Figure 2D).

There is an ongoing lack of clarity regarding the levels of DNA methylation that determine change in DNA activity during human folliculogenesis. Here we demonstrated high expression levels of *DNMT1*, *DNMT3A* and *DNMT3B* in oocytes at all stages of folliculogenesis with mounting abundance (Figure S2A), which implies that DNA methylation level might be continually elevated with oocyte maturation. In contrast to our previous study in PGCs ^16^, our results showed that ten-eleven translocation (TET) family genes were under-expressed in oocytes at all stages of development (Figure S2A). Together, overexpression of DNMT genes with loss of TET activity indicates an active methylation transition in human oocyte during folliculogenesis.

### Dynamic Expression and Transcriptional profiles of GCs during Folliculogenesi

In GCs, PCA of all samples showed clustering into five groups according to stage of follicular development (Figure 3A). An observed overlap between the clusters of primary and secondary follicles indicated less pronounced difference at the transcriptome level between these two stages. We then examined DEGs (FDR < 0.05; FC of log_2_ transformed FPKM > 1.5) in GCs (Figure 3B). Interestingly, some of the DEGs have been previously reported in association with oocyte competence, including quality, maturity or fertilization rate, embryo quality and pregnancy outcomes ^18, 19^ as detailed in Table S2. GO analysis of DEGs (Figure 3C) revealed the crucial functional roles of these genes during folliculogenesis. The expression profiles of the DEGs *CDCA3*, *BNIPL* and *TST* were validated by using immunohistochemistry (IHC) on ovarian tissue from the scRNA-Seq sample set (Figure 3E), which are in concordance with the scRNA-Seq results.

**Figure 3.**
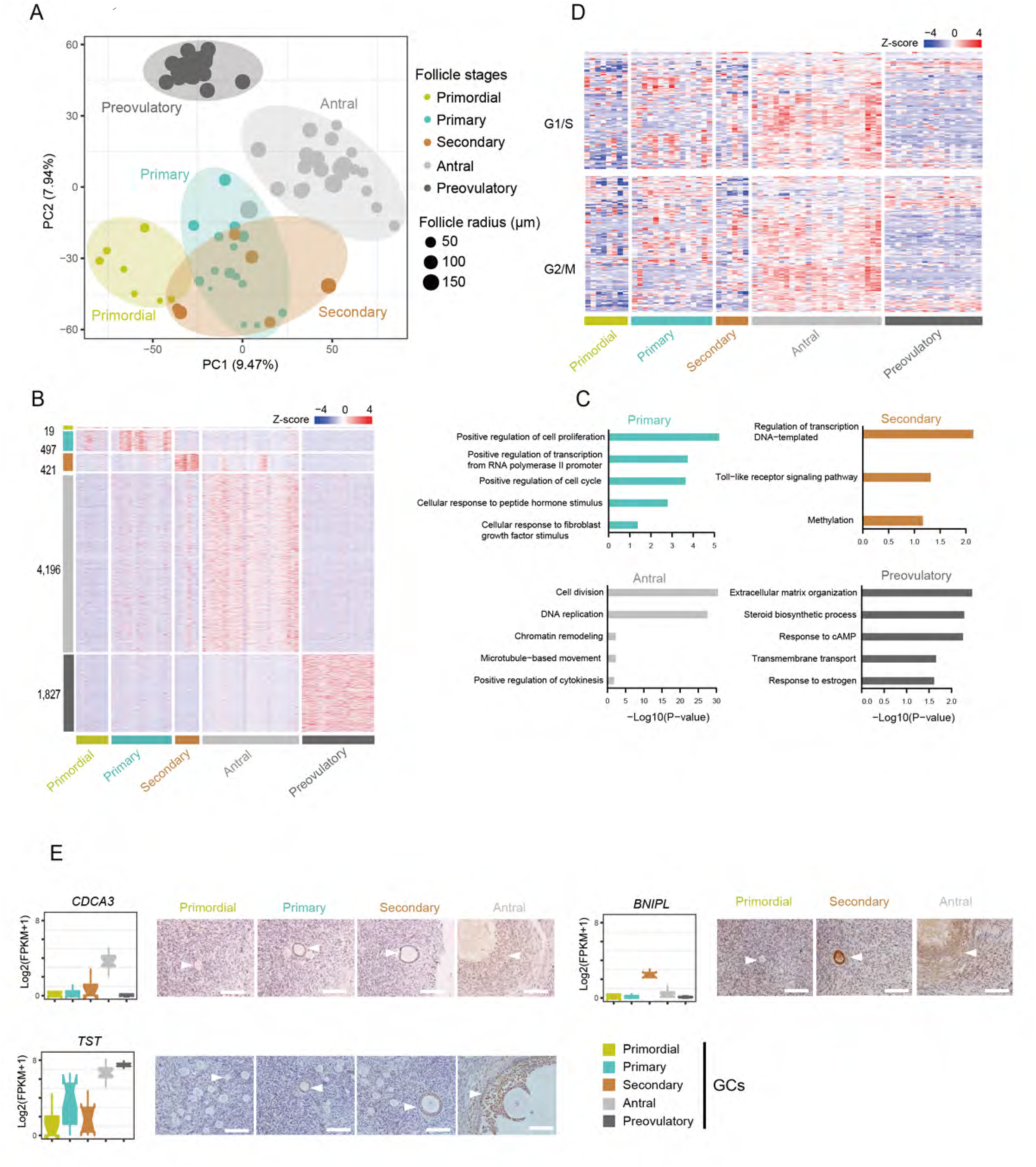
Dynamic Gene Expression Patterns and Transcriptional Features in GCs. A) Principal component analysis (PCA) of the transcriptome of scRNA-Seq data from 71 GCs collected from follicles at different stages of development. GCs are clustered into five subpopulations. Different colors and sizes of the points represent follicles of different stages and sizes, respectively. B) Heatmap of all the differentially expressed genes (DEGs) in GCs at five stages of folliculogenesis. The numbers of identified DEGs are indicated on the y-axis, and the stages of follicle development are presented along the x-axis. The color key from blue to red indicates the relative gene expression level from low to high, respectively. C) Significantly enriched GO terms (biological processes) of DEGs in GCs at five stages of folliculogenesis. D) Heatmap of cell-cycle related genes that are expressed in GCs at five stages of folliculogenesis. E) Immunohistochemistry staining of selected DEGs. The boxplot demonstrating gene expression level is presented on the left of each corresponding immunohistochemistry panel. White triangles indicate the follicles. Scale bar, 100 µm.

Analysis of the cell-cycle genes involved in the G1/M and G2/S phases ^20^ in GCs Revealed that at primordial stage, GCs were found to be relatively quiescent and maintained low proliferative activity (Figure 3D). Both G1/S and G2/M-specific genes were up-regulated in the GCs from primary, secondary and antral follicles. The cell-cycle genes were predominantly abundantly expressed in the GCs from antral follicles, which designates their high proliferative activity at this stage. In contrast, the cell-cycle genes were down-regulated in the GCs from preovulatory follicles, which implies decreased degree of proliferation and differentiation in GCs at this stage.

Next, we focused on genes that have been previously associated with steroidogenesis (Figure S2B). *CYP11A1*, *CYP19A1*, *HSD3B1* and *HSD17B2*, encoding for steroidogenic enzymes, showed up-regulation in the antral follicles and reached peak expression levels in the preovulatory follicles. Their expression patterns indicate progressive increase in steroidogenic activity in maturing follicle, which culminates before ovulation. *NR5A1*, a key regulator of steroidogenic enzyme-encoding genes that has been previously described in sheep GCs ^21^, exhibited up-regulation with the progression of follicle growth. Both *ESR1* and *AR*, encoding for estradiol (E2) and androgen hormone receptors, respectively, were expressed in GCs at the primary follicle stage and reached highest expression level in antral follicles. This suggests E2 and androgens might control GC growth from the primordial follicle stage through these receptors.

### Stage-Specific Signature Genes Identified in Oocytes and GCs

We identified a subset of stage-specific oocyte genes that were exclusively pertained to a single stage of folliculogenesis, which we referred to as “signature genes”. Conjointly, 382 signature genes were identified comprising the genes expressed in the oocytes from each stage (Figure S3A, Table S3). Distinct stage-specific expression patterns of *NTF4* and *LCP2* in oocytes were further confirmed by IHC staining (Figure 2E), which showed that the expressions of these genes were restricted in antral stage. Of note, *NTF4*, proposed to facilitate follicle development in mouse by inducing FSH receptor (*FSHR*) ^22^, was exclusively expressed in oocytes from antral follicles (Figure 2E). We found that *FSHR* gene expression was pertained to GCs in antral follicles and was concordant to that of *NTF4* (Figure S3D). Our data suggested that *NTF4* might upregulates *FSHR* expression in human GCs and contributes to oocyte-to-GC crosstalk, thus plays an important role in oocyte development. For GCs, we also identified a subset of stage-specific signature genes during folliculogenesis (Figures S3C and S3D, Table S3). The expressions of signature genes *CDCA3* and *BNIPL* were validated by IHC (Figure 3E). Interestingly, *FSHR*, previously observed in somatic cells from early preantral to mature follicles in sheep ^23^, was exclusively expressed in the GCs from antral follicles in our study (Figure S3D), suggesting considerable inter-species differences. The full list of the signature genes in oocytes and GCs is presented in Table S3. These identified signature genes (Figure S3 and Table S3) could serve as candidate cell-specific markers for each follicle stage, allowing to establish the objective criteria for selecting competent oocytes in vitro.

### Secretory Protein Coding Signature Genes

Certain specific oocyte- or neighboring GCs-derived genes could reflect the quantity and the quality of the follicles. Consequently, the peripherally secreted products of these genes may reveal prognostic information concerning ovarian reserve. Therefore, we searched for the secretory protein-coding signature genes that were exclusively pertained to either oocytes or GCs and were expressed either at early or at more advanced stages of follicle development (as described in Methods). In total, we identified 51 protein-coding genes in oocytes and 17 in GCs that clustered into five groups (Figure S4). Cluster 1 comprised 16 oocyte genes with specific expression in the oocytes of primordial and primary follicles, suggesting they may reflect the primordial follicle pool. Cluster 2 included 19 oocyte genes with specific expression in the oocytes from secondary, antral and preovulatory follicles. Cluster 3 consisted of 27 oocyte genes that were represented across folliculogenesis. Cluster 4 (n = 4) and cluster 5 (n = 13) included GC-derived genes with specific expression at preovulatory and secondary-antral stages, respectively. The expression patterns of the genes from each cluster are exemplified in Figure S4B. These findings represent a potentially valuable source of candidate molecular biomarkers of ovarian reserve. Further evaluation of their expressions in biological fluids could contribute to development of ovarian reserve test.

### Transcription Factors Regulatory Networks in Oocytes and GCs

To find the master regulators and construct the transcriptional regulatory network along the steps of human folliculogenesis, we utilized the ARACNe method to analyze all 1,469 known TFs from Animal Transcription Factor Database (TFDB v2.0) ^24, 25^. In the oocytes, the expressions of *GTF2I, CSDE1, SOHLH2, SMARCE1, TUB, HBP1, SOX30* and *HIF1A* were up-regulated in primary follicles, indicating that these TFs may play a critical role in the transition from primordial to primary stage (Figure 4). *KLF2, YBX2, FOXO6, SOX13, ETV5, TEAD2* and *OTX2* were over-expressed in the oocytes from secondary follicles compared to those from primary follicles, implying they are likely the regulators of primary-to-secondary stage transition. *PINX1, PBX1, MTF1, SOX15, UBTF, SOX13,* and *POU2F1* had higher expressions in the oocytes from antral compared to secondary follicles, indicating their crucial regulatory roles in cytoplasmic and nuclear maturation of oocyte during antral stage. *ATF2* and *EOMES* were abundant in MII oocytes of preovulatory follicles, which indicates their potential role as TFs that possibly initiate the unique transcription networks involved in meiosis progression. Of note, we found that *SOX13* and *SOX15,* members of the SOX family of TFs, were up-regulated in the oocytes from secondary to antral follicles. While *SOX30* was found abundantly expressed in the oocytes during transition from primordial to primary follicle stages. This indicates their potential role in regulating transcription networks during primordial follicle activation (*SOX30*) and antral formation (*SOX13, SOX15*), respectively.

**Figure 4.**
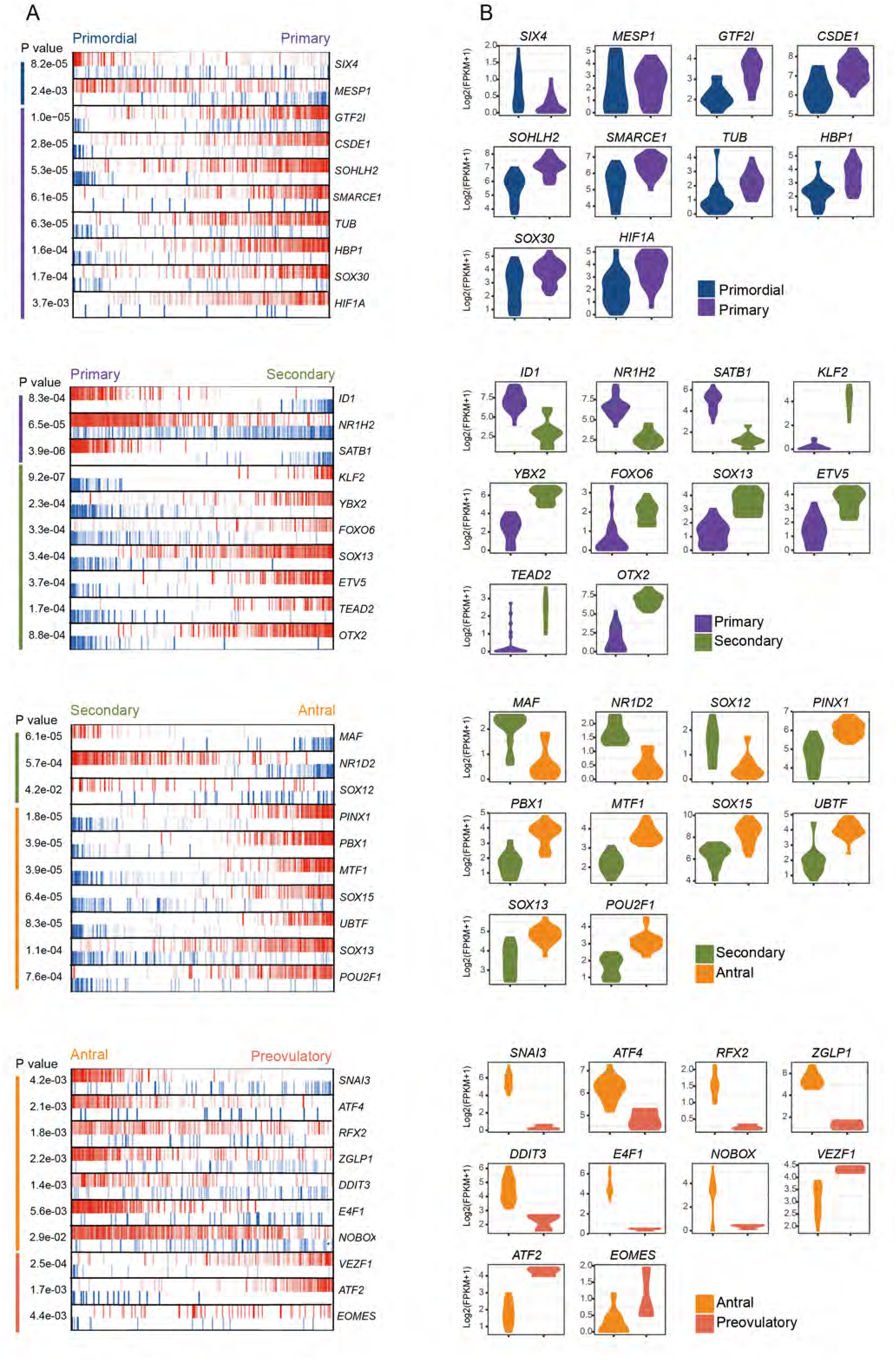
Inferred Key Transcriptional Factors in Oocytes at Each Stage-to-stage Transition of Folliculogenesis. A) MARINa plots of targets for each candidate master regulator. Red vertical bar represents the activated targets, blue represents the repressed targets. On the x axis, genes are rank-sorted according to the significance of differential expression between the two developmental stages. The candidate master regulators are displayed on the right. The corresponding p-values of these master regulators indicate the significance of enrichment are displayed on the left. See also Figure S4 for the analogous information in GCs. B) Violin plots show the relative expression levels (log2 [FPKM+1]) of each master regulator in two consecutive stages. See also Figure S4 for the analogous information in GCs.

In the GCs, we found that *CREB1, NFKB1, MEF2A, PIAS1, FOSL2, KLF13* and *PRDM4* potentially regulated activation from the primordial to primary follicle stage, whereas *CRX, HES2, ZNF554, ZKSCAN3, LMX1B, FOXK1* and *SIX4* were the top candidate TFs possibly involved in transition from the primary to secondary follicle stage. *MBD1, FIZ1, GABPA, TGIF2, PIAS3, E4F1* and *IRF3* were likely driving initiation of the transcription network in antral follicles. *MEIS3, PRDM15* and *VTN* expressed in GCs of preovulatory follicles, could be the key regulators of cumulus cells progression in preovulatory follicles (Figure S5). Further evaluation of these TFs, their upstream signaling pathways, ligands, receptors and downstream targets in oocytes and GCs will provide insight into the transcriptional control of human folliculogenesis.

### Key Signaling Pathways in Oocytes and GCs during Folliculogenesis

The Gene Set Enrichment Analysis (GSEA) and Kyoto Encyclopedia of Genes and Genomes (KEGG) analysis were applied to perform the pairwise comparisons of the follicular stages associated with each transition. There were 43 pathways overrepresented in the oocytes and 79 pathways enriched in the GCs from the primary follicles versus the primordial follicles (*p* < 0.05) (Table S4). Of these, the functional pathways that were significantly overrepresented both in oocytes and GCs (*p* < 0.05) and described in a context of folliculogenesis, included mTOR, GnRH, Neurotrophin and Insulin signaling (Figures S6A and S6B). These pathways probably mediate primordial-primary follicle transition and indicate the concerted action of the two follicular compartments. We identified 21 pathways enriched in the oocytes and 60 pathways in GCs that were overrepresented in antral, but not in secondary follicles (*p* < 0.05) (Table S4). In GCs, the steroid hormone biosynthesis was among the enriched pathways at the secondary-to-antral follicle transition, which indicates GCs play a key role in steroidogenesis at antral stage ^26^. In addition, mTOR, Insulin and Neurotrophin pathways were over-represented in both primordial-to-primary and secondary-to-antral stages, suggesting a possibility for their involvement in these two transitions (Figures S6B, S6C and S6D). Interestingly, the enrichment of Neurotrophin pathway was concordant with the expression of its ligand *NTF4*, which was identified in this study as an antral stage-specific signature gene in oocyte (Figure 2E).

### Bidirectional Interactions between the Oocyte and GC Compartments

To investigate interactions between oocyte and GCs, first we analyzed the expression of the components of cell signaling pathways, including ligands, receptors and target genes. NOTCH pathway was one of the significantly enriched in antral follicle GCs (Table S4). Expression of the components of NOTCH signaling revealed that ligands *DLL3* and *JAG1* were predominantly expressed in early oocytes before preovulatory stage. Their receptors *NOTCH2*, *NOTCH3* and the downstream target gene *HES1* were highly expressed in GCs (Figure 5A). This finding highlights the role of NOTCH signaling in oocyte-controlled proliferation and differentiation of GCs.

**Figure 5.**
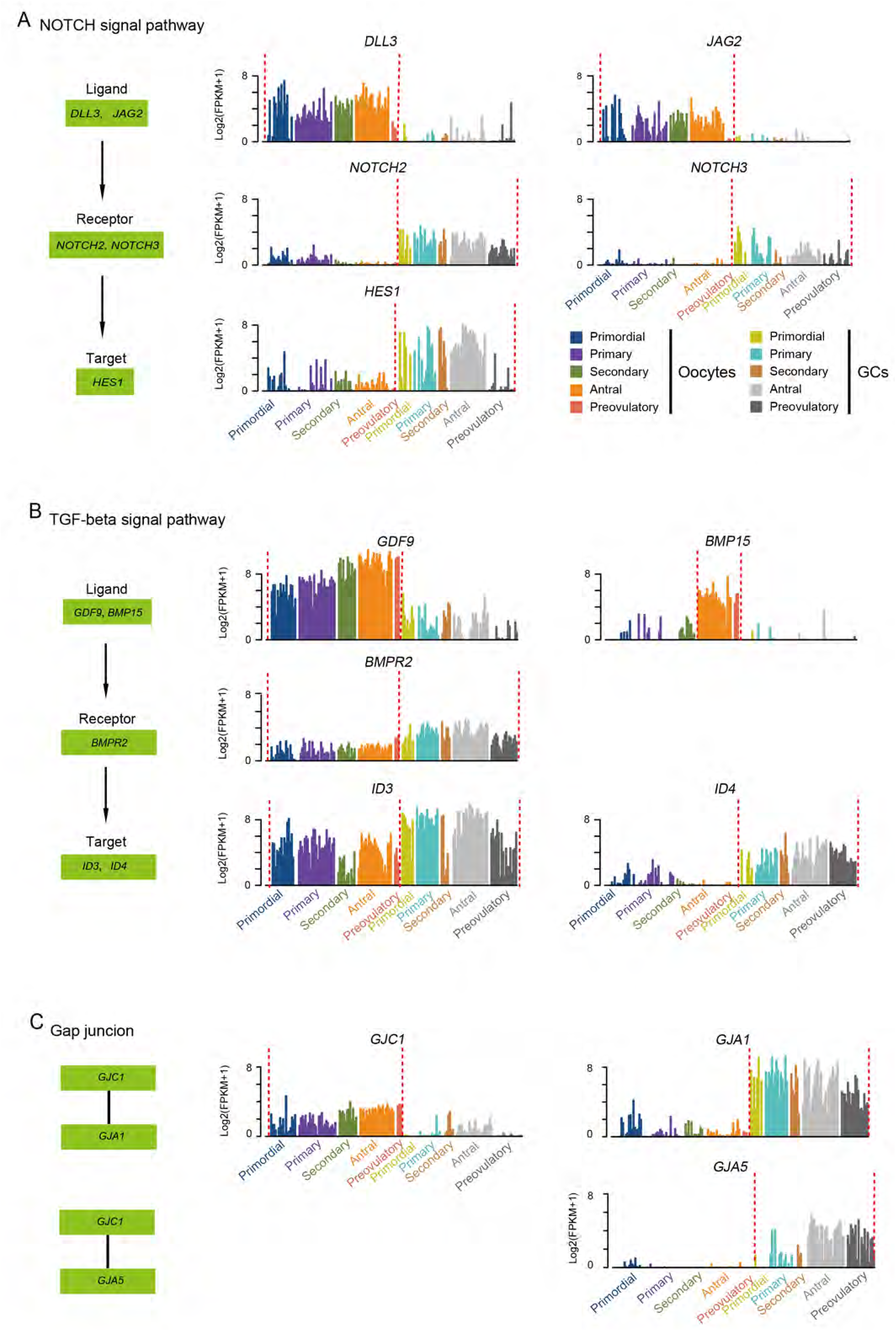
Signaling Pathways and Gap Junction Involved in Oocyte-GC Crosstalk. A) NOTCH signaling pathway involved in oocyte-GC crosstalk in follicogenesis. The relative expression levels (log2 [FPKM+1]) of the specific ligands, receptors, and target genes are shown. The diagrams at the left show the relationship among these genes. B) TGF-β signaling pathway involved in oocyte-GC crosstalk. C) Gap junction involved in oocyte-GC crosstalk. The relative expression levels (log2 [FPKM+1]) of the connexin genes are shown. The diagrams at the left show the putative relationship among these genes.

Then we analyzed the expression of the key components of TGF-β signaling pathway in oocytes and GCs. Our data revealed that the expression levels of *GDF9* in oocytes were high at all stages of follicular development, whereas *BMP15* was highly expressed only in the oocytes from antral and preovulatory follicles (Figure 5B). This observation points out that *GDF9* is likely to activate the TGF-β all through folliculogenesis, while *BMP15* mostly exerts its role at more advanced stages. In contrast, animal studies demonstrated that activation of the primordial follicles is mediated by *BMP15* ^27^, suggesting it might play different roles in folliculogenesis across the species.Interestingly, BMP type II receptor gene *BMPR2* and its target *ID3* were expressed in both oocytes and GCs. The presence of gene and its target in two different cell types, herein, allows to assume an existence of both autocrine and paracrine mechanisms in BMP signaling.

In addition, we identified the components of GC-derived signaling involved in folliculogenesis. For example, *KITLG* and its receptor *KIT,* previously implicated in paracrine signaling in folliculogenesis, were expressed in GCs and oocytes, respectively, which is consistent with previous findings in animal models ^28^. *KITLG* was expressed in primordial and upregulated in primary follicles with a subsequent down-regulation, suggesting a possible role in primordial follicle activation (Figure S2). Together, our findings provide evidence that human folliculogenesis is coordinated by both autocrine and paracrine signaling pathway mechanisms that could be initiated by either oocytes or GCs.

To understand the distribution of gap junctions in human follicles, we screened both oocytes and GCs for the connexin (gap junction components) encoding genes, previously reported in mammalian ovary, namely *GJB2* (Cx26), *GJB4* (Cx30.3), *GJB1* (Cx32), *GJA4* (Cx37), *GJA5* (Cx40), *GJA1* (Cx43), *GJC1* (Cx45) and *GJA10* (Cx57) ^29^. Three out of these genes exhibited compartment-specific pattern across the follicle stages, while others were largely under-expressed. We identified that *GJC1* (Cx45) was expressed only in the oocytes and *GJA1* (Cx43) was exclusively expressed in the GCs at all follicle stages, while *GJA5* (Cx40) was pertained to the GCs of antral and preovulatory follicles (Figure 5C). We speculate that the heterotypic (Cx43/Cx45 or Cx40/Cx45) connexins might generate gap junction channels between oocyte and GCs, while homotypic (Cx43 or Cx40) connexins may generate the channels between GCs.

### Global Expression Patterns of Maternal Effect Genes

To explore the global expression profiles of maternal-effect genes, we integrated the oocyte transcriptome data derived from this study with the transcriptome of human preimplantation embryos that had been identified in our previous work ^30^. We searched for the genes that were expressed in MII oocytes and in zygotes, but not expressed in 8-cell embryos, as these genes are carried over from oocyte to early embryo and then get degraded following zygotic genome activation (ZGA). Overall, we identified 1,785 putative maternal genes. The unsupervised cluster analysis identified 4 clusters of genes according to expression patterns at specific stages of follicular growth (Figure S7A, Table S5). The genes in cluster 1 (n = 47) and cluster 2 (n = 588) were not expressed in primordial stage and upregulated during advanced stages of follicular maturation with peak expression in MII oocytes. However, the expression patterns in cluster 3 (n = 987) and cluster 4 (n = 163) showed that these transcripts could accumulate before primordial follicle assembly and potentially could be involved in early folliculogenesis and embryo development. The GO analysis revealed biological functions that have been reported in association with meiosis were among the overrepresented terms in cluster 2. While in cluster 3 and 4, genes were enriched in ubiquitous functions (Figure S7B).

### Comparison of Human and Mouse Oocyte Transcriptomic Profiles during Folliculogenesis

Appraisal of the inter-species differences in oocyte transcriptome dynamics during all the stages of folliculogenesis is important to better understand the translational limitations of animal oocyte models, but such comparative data are largely lacking. Recent meta-analysis demonstrated substantial variation in oocyte transcriptome across different species ^31^. To better understand the interspecies complexity of oocyte transcriptome in human and mice, and to provide a comparative view of gene expression in human and mice oocytes across folliculogenesis, we compared our findings in human oocytes with reported RNA-Seq datasets produced from mice oocytes collected from primordial to antral stage. Mice RNA-Seq data were downloaded from Gahurova’s study^32^. The standardized analytical approach to the merged raw data revealed a total of 23,091 genes expressed in human (our data) and 11,585 genes expressed in mice oocytes (Figure 6A).

**Figure 6.**
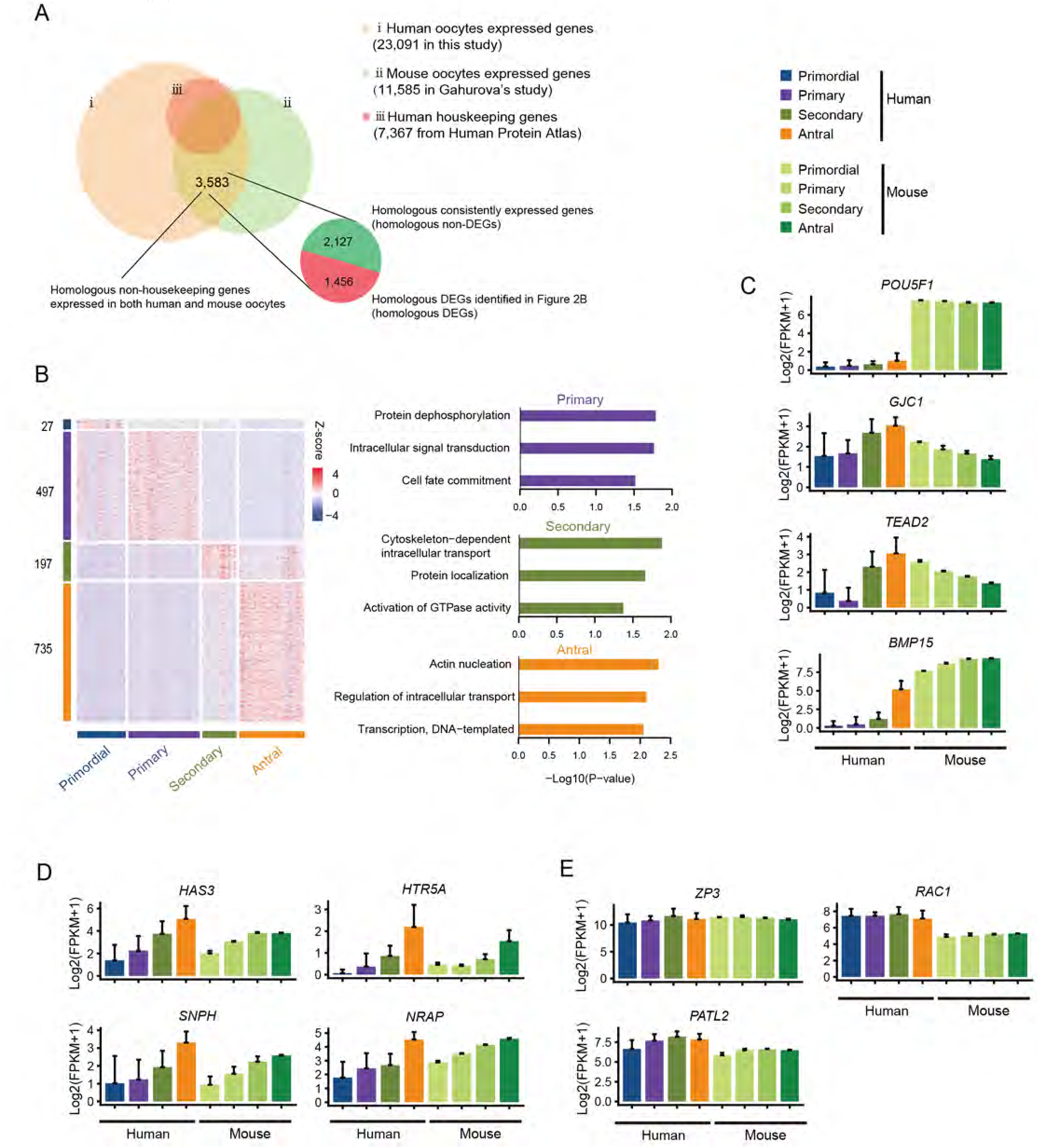
Comparison of Human and Mouse Oocyte Transcriptomic Profiles during Folliculogenesis. A) Venn diagram of total expressed genes in human oocytes (peach) and mouse oocytes (light green) demonstrates overlap between the gene populations in two species and also overlap with the human housekeeping genes (deep orange). Pie chart represents the non-housekeeping human-mouse oocyte-derived homologous genes (3,583), including homologous DEGs (red) and homologous ubiquitous genes (green). B) Heatmap of homologous DEGs in oocytes at five stages of folliculogenesis. The numbers of identified DEGs are indicated on the y-axis, and the stages of follicle development are presented along the x-axis. The color key from blue to red indicates the relative gene expression level from low to high, respectively. The enriched GO terms of human oocytes are shown on the right. C) Bar plots demonstrate a comparative analysis of the selected oocyte-derived homologous DEGs that show the distinct expression patterns in human (left) and mouse (right) oocytes during folliculogenesis. Different colors represent different stages of folliculogenesis. D) Bar plots demonstrate a comparative analysis of four oocyte-derived homologous DEGs that show the similar expression pattern in human (left) and mouse (right) oocytes during folliculogenesis. E) Bar plots demonstrate a comparative analysis of three homologous non-DEGs annotated as oocyte maturation defect genes (OMIM) in human (left) and mouse (right) oocytes during folliculogenesis.

We focused further analyses on 16,175 one-to-one homologous genes shared by human and mice, as per Mouse Genome Informatics (MGI) database ^33^, of which 14,431 genes were expressed in human oocytes and 9,964 genes were expressed in mice oocytes. There were 9,698 homologous genes that overlapped between human and mice oocyte samples. Of these, there were 6,115 house-keeping genes, 2,127 consistently expressed genes and 1,456 DEGs identified in this study as differentially expressed between different stages of folliculogenesis (Figures 6A). GO analysis of the co-expressed DEGs revealed that these genes exhibited ubiquitous functions, and none were specific for oocyte development (Figure 6B), which suggested that distinct molecular mechanisms may be involved during folliculogenesis in human and mouse. Moreover, several homologous DEGs implicated in oocyte development, such as *POU5F1,GJC1, TEAD2* and *BMP15*, showed different expression patterns in human and mice oocytes (Figure 6C).

Next, we assumed that oocyte genes with concordant expression pattern and high degree of correlation (r > 0.8) are involved in conserved molecular mechanisms in human and mice oocytes (Figure 6D). Notably, consistent homologous genes (*ZP3*, *RAC1*, *PATL2*), annotated in association with “oocyte maturation defect” in Online Mendelian Inheritance in Man (OMIM) database, showed stable expression across folliculogenesis in human and mice oocytes (Figure 6E), which proposes conserved functional importance of these genes for oocyte development in both species. These shared expression patterns suggested existence of evolutionary conserved mechanisms involved in folliculogenesis and highlights the potentially significant translational value of these conserved genes to the future research.

## Discussion

This work describes the transcriptional landscape of the human oocytes and their surrounding GCs across all major stages of folliculogenesis. To the best of our knowledge, this is the first comprehensive investigation of human transcriptome from both the germ cell and somatic follicle compartments in the adult ovary at single-cell resolution. Previously, we have characterized the transcriptome profiles of human pre-implantation embryos and PGCs with their gonadal niche at several stages of development ^16, 17, 30^. It is well established that oogenesis, folliculogenesis and embryogenesis are sequential interrelated events, thus the transcriptome features of both fetal oogenesis and adult folliculogenesis confer the underlying molecular mechanisms of oocyte development. This body of work complements the findings from our previous studies and contributes to a comprehensive overview of the transcriptional events and their dynamics throughout the course of female gametogenesis with a subsequent link to early pre-implantation embryo.

Previously, characterization of ovarian follicle in humans relied on evaluations of the oocytes at certain stages of development or of the unseparated follicle compartments ^12–14^. Although previous studies provided important insights into molecular events involved in folliculogenesis, generalizing the conclusions is made challenging by the multiple technical aspects related to different testing platforms and analytical variables. Tissue heterogeneity is one of the main predicaments of biological research when it is crucial to distinguish cell-specific profiles that reflect the in-vivo status. Laser Capture Microdissection (LCM) is increasingly used to harvest the cells of interest and to improve experimental precision, although faces challenges of compromised RNA quality and cell integrity ^34^. To date, there have been no other studies that evaluated the transcriptomic profile across all the key stages of folliculogenesis and that performed parallel evaluations in both oocytes and GCs.

For the first time we demonstrated that each stage of folliculogenesis, defined by the morphological characteristics of both follicle compartments, had distinct transcriptome profile with different gene expression dynamics in oocytes and GCs. This finding supports the proposed, but not entirely proven concept that morphological classification of ovarian follicles reflects complex intrinsic transcriptional changes during folliculogenesis.

We also identified the candidate compartment-specific and stage-specific genes in both oocytes and GCs that provide a valuable clue for future more directed functional research. These findings may have important implication for development of essential genetic tools for cell-type-specific or stage-specific labeling and manipulations, which could be utilized in human follicle functional studies ^35^. Further, an unmet need for the reliable predictors of oocyte developmental competency drives discovery of new biomarkers that would allow to establish more objective criteria for selecting competent oocytes ^36^. Previously reported biomarkers associated with successful embryo development and pregnancy outcomes were mainly derived from GCs and showed little consistency across the studies, which could be explained by the differences in experimental methodology ^18, 19^. Some of the already proposed biomarkers exhibited distinct expression patterns in our study, which makes them particularly attractive candidates for further investigations. Of note, the functional characteristics for majority of the gene markers identified in this study are unrecognized and await further investigation.

In addition, we identified a subset of regulators of oocyte and GC transcriptome that may be involved in follicle development and acquisition of oocyte competency. Further evaluation of these TFs, their upstream signaling pathways, ligands, receptors and downstream targets in oocytes and GCs will provide more understanding of the transcriptional control of human folliculogenesis. These TFs may evolve as potential targets for future therapeutic interventions to modulate follicle development.

It has been widely acknowledged that oocyte-granulosa bidirectional communications are one of the core mechanisms involved in oocyte acquisition of developmental competence ^4^. Communication via signal transduction (e.g. signaling pathways) and via direct cell-to-cell contact (e.g. gap junctions) provide molecular and structural basis for effective oocyte-GC crosstalk required for follicular development. However, precisely how oocyte-GC interact and how bi-directional communications between the follicle compartments are initiated and maintained, is still unclear. In this study we uncovered the signaling pathways that are coordinately and reciprocally regulated in human oocytes and their surrounding GCs, by identifying the ligands and their receptors that were derived from the reciprocal compartments, but shared the expression pattern. This indicates that signaling pathways are activated by the ligands derived from the germ cells that act on the neighboring gonadal somatic cells, and vice versa. For example, in this study NOTCH signaling pathway, known important regulator of cell-cell communication ^37^, was shown to be activated in GCs via oocyte-driven mechanisms, which is in line with previously observed by our group in fetal ovary ^17^. In contrast, KITLG-KIT pathway was GC-driven, while TGF-β pathway appeared to be activated through both autocrine and paracrine regulations. Single-cell resolution made possible evaluation of the distinct gap junctional channels in human follicles. We identified three different connexins, known to contribute to gap junctions in animals, and we infer their involvement in gap junctions in humans, which so far has been only demonstrated in animal models. Together, our findings provide further insight into molecular basis of bidirectional interactions in both cell signaling pathway and gap junction communications in human follicle. We support the notion that human folliculogenesis is coordinated by both autocrine and paracrine signaling pathway mechanisms that could be initiated by either oocytes or GCs.

In this study, we propose a set of candidate biomarkers that could be a valuable source of information on reproductive potential and ovarian reserve. Ideally, an ovarian reserve test should be easy to perform, reproducible and, importantly, should clearly show evidence of improvements in patient reproductive outcomes following test-directed interventions. However, none of the currently available ovarian reserve tests, namely anti-Mullerian hormone (AMH) ^38, 39^, follicle stimulating hormone (FSH) and the antral follicle count (AFC) measured by transvaginal ultrasound ^40, 41^ seem to meet the above criteria. It is now understood that while ovarian reserve markers can moderately predict response to stimulation in assisted reproductive technology (ART) cycles, none of these tests determine the pregnancy potential and their role in general population or in infertile women not undergoing ART treatment is not confirmed ^10, 42^. For instance, AMH, originating from the GCs of secondary and antral follicles, has been initially accepted as the endocrine marker of FOR in humans ^43^. However, it has been recently proposed that AMH is not associated with the reproductive potential in women of late reproductive age ^8^ and is a moderate predictor of menopause in general population ^9^. The main limitation of the current approach is inability to directly measure ovarian reserve, since the biomarkers to determine the number or quality of primordial follicles, a non-renewable pool that determines reproductive life span ^44^, have yet to be identified. In addition, majority of the available biomarkers are secreted by GCs and currently there is little information on oocyte-derived markers that provide information on a status of follicle pool. Furthermore, it is possible to assume that ‘all-purpose biomarker’ does not exist, and different markers would be required to address the specific outcome of interest. For example, reproductive potential in general population is likely to be represented by the markers expressed at all stages of follicle development, while age of menopause could be more accurately predicted by the markers that originate in primordial pool. It is also possible that response to fertility treatments and pregnancy outcomes may be better predicted by the markers confined to the antral follicles, the main targets of ovarian stimulation gonadotropin treatments, while the predictors of natural pregnancy outcomes may be secreted by the more advanced follicles. Until new data emerge, these questions will remain open. Further research to elucidate the biological roles of the proposed candidates, in conjunction with comprehensive evaluation of their expression levels in biological fluids could contribute to advancements in development of clinically informative ovarian reserve test.

In this study we offer further insights on species-specific gene expression during folliculogenesis. Mouse models are a mainstay of translational biomedical research and are widely implicated in the investigations of ovarian transcriptome. Mice are poly-ovulating species, breed readily, inexpensive and easy to handle, which makes them ideal candidates for laboratory experiments. Besides, mice and humans are at least 95% genetically identical, which allows to assume conservation of fundamental biological processes across the species. However, it has been previously reported that the oocyte transcriptome is highly variable across mammals and the human oocyte is likely to have a greater complexity than other mammals ^31, 45^. Appraisal of the inter-species differences in oocyte transcriptome dynamics during all the stages of folliculogenesis is important to better understand the translational limitations of animal oocyte models, but such comparative data are largely lacking. We evidenced strong transcriptional activity in both human and mouse oocytes, which dominated in humans and demonstrated considerable variability in oocyte transcriptome between human and mouse. We revealed different gene expression dynamics during folliculogenesis in both species and thus, propose a cautious approach when mice oocyte data are applied to the human domain. We also showed some degree of similarity in gene expression between human and mice oocytes, which highlights the potentially significant translational value of these conserved genes to the future research. We do not suggest that the mouse is an invalid experimental system for studying human oocytes. Rather, we provide a database for further investigations of the molecular mechanisms associated with oocyte development through mouse models, and improve the understanding on how well the oocyte transcriptome data translate from mouse to human.

It is important to mention that our study is not devoid of limitations. Firstly, despite rigorous experimental approach and robust analyses, our findings are not sufficient to confirm the functional characteristics of the identified transcripts. Rather, our data offer a comprehensive resource for future more directed functional studies in oocyte research. Next, our methods relied on the mechanical or enzymatic dissociation of follicular compartments, thus the information on temporal and spatial regulation of gene expression in GCs was not reflected. The recently proposed Geo-Seq protocol allows the profiling of transcriptome information from only a small number cells and retains their native spatial information ^34^, which will help in future investigations of gene expression with positional information. Finally, low number of MII oocytes included in this study could undermine the reliability of the results for this group, although satisfactory gene number and considerable homogeneity of the expression data from these cells suggest adequate quality of the data.

In summary, understanding molecular events involved in folliculogenesis remains a key challenge for reproductive biology research with limited availability of human follicles for research being the main constrain. By using a high-resolution transcriptome analysis for single oocyte and its surrounding GCs, we recapitulated a cascade of the molecular events involved in folliculogenesis. This work contributes to the collaborative effort to strengthen our understanding of reproductive function, which may assist with the development of more targeted future interventions to improve oocyte competence in-vitro and in-vivo.

## METHODS

### Collection of Human Ovarian Tissue Samples

With oral and written informed consent, fresh ovarian tissues were taken from 7 female donors who underwent ovariectomy for the following indications: sex reassignment surgery (n = 1), fertility preservation for cervical cancer (n = 1), endometrial cancer (n = 2), benign ovarian mass (n = 2) and lymphoma (n = 1). All the donors were of reproductive age, ranging 24 to 32 years (median age of 28 years). All the participants had regular menstrual cycles, had no history of autoimmune or genetic conditions. None of the tissue donors were on hormonal treatment at least 6 months before surgery, had no previous ovarian surgery and were not exposed to any cytotoxic agents or radiotherapy. All the samples had normal histopathology (described below). The ovarian tissue samples were collected in operation theatre during the procedure, immediately transferred to the laboratory and treated as previously described ^46^. Briefly, the human ovarian tissues were collected in the operation theatre and transported to the laboratory in Leibovitz’s L-15 medium (Sigma-Aldrich) on ice with the supplementation of 1% human serum albumin (Life Technologies, Carlsbad, CA), 100 IU/mL streptomycin (Sigma, St. Louis, MO) and 100 µg/mL penicillin (Sigma, St. Louis, MO). Scalpel and surgical scissors were used to enucleate the medulla tissues. Then scalpel was used to cut the ovarian cortical tissues into small ovarian pieces with a size of 5 mm×5 mm×1 mm.

### Ovarian Histology Assessment

Histological assessment was performed on all the ovarian tissue samples by using hematoxylin and eosin (HE) staining as described elsewhere ^47^. After fixation, the samples were paraffin embedded and cut into serial sections of 5-µm-thick. All the prepared tissue sections were reviewed by two independent pathologists and confirmed normal ovarian tissues.

### Immunohistochemistry

Immunohistochemistry staining of the ovarian tissue samples that were utilized in this study for RNA-Seq experiments was performed by using the ABC Staining System (Zhongshan Golden Bridge Biotechnology, Inc., Beijing, China), as previously described ^48^. Brown staining of the cytoplasm or nucleus of the cells was considered as positive.

### Isolation of Human Oocytes and GCs

Human follicles were isolated from fresh ovarian tissues as described previously ^49, 50^. Briefly, after removal of medulla tissues, the ovarian cortical pieces were placed in a tissue sectioner (McIlwain Tissue Chopper, The Mickle Laboratory, Guildford, UK) and cut into 0.5 × 0.5 × 1 mm pieces. Then the tissue pieces were enzymatically digested by a mixed digestion medium, which included αMEM (Sigma-Aldrich) media, 0.04 mg/ml Liberase DH (Dispase High; Roche Diagnostics GmbH, Mannheim, Germany), 10 IU/ml DNase I (Sigma-Aldrich), 100 IU/ml penicillin and 100 µg/ml streptomycin (Invitrogen) and incubated for 75 min on a shaker at 37°C (Thermo Fisher, Marietta, OH, USA). The incubation was terminated by double wash with DPBS (Sigma-Aldrich) supplemented with 10% HSA (LifeGlobal). Then the follicles were isolated mechanically using 29G needles and were transferred to culture medium. Subsequently, the single oocytes and GCs were enzymatically separated with accutase (Sigma-Aldrich) and were mechanically isolated by using 29G needles. GC samples comprised randomly selected 10 cells per sample because of low abundancy of RNA in these cells. GCs from the antral and preovulatory follicles included cumulus cells isolated from the cumulus-oocyte complex (COC) as following: 6 samples of cumulus cells from the antral follicles and 16 samples from the preovulatory follicles. Follicle stages were classified according to the criteria described by Gougeon ^51^. The diameters of follicles and oocytes were measured under light microscope (Nikon).

### Single-Cell cDNA Libraries Construction from Oocytes and GCs

The oocytes and GCs that were isolated from follicles were analyzed by scRNA-Seq as previously described ^30, 52^. Briefly, the single oocytes or 10 randomly selected GCs were transferred into the lysis buffer quickly using a mouth pipette. Then we performed reverse transcription on the cell lysate and terminal deoxynucleotidyl transferase was adopted to add a poly A tail to the 3′ end of the first-strand cDNAs, next we performed 20 cycles of PCR to amplify the single-cell cDNA library. qPCR analysis was conducted to check the quality of the cDNA libraries using housekeeping genes (GAPDH and RPS24). The RNA-Seq libraries were constructed by a Kappa Hyper Prep Kit (Kappa Biosystems).

### RNA-Seq Data Processing

The analysis of single-cell RNA-Seq data was carried out as previously described ^16, 17, 53^. Briefly, RNA-Seq raw reads with 10% low quality bases, adapters and artificial sequences (including UP1, UP2, polyA sequences) introduced during the experimental processes were trimmed by in house scripts. Next, the trimmed clean reads were aligned to the UCSC human hg19 reference using Tophat2 (v2.1.0) with default settings ^54^. Cufflinks (v2.2.1) was further used to quantify transcription levels of annotated genes ^55^.

Previous published data, including those from human MII oocytes ^30^, human pre-implantation embryos ^30^, and mouse oocytes ^56, 57^, were downloaded from the GEO data sets, and the raw fastq reads were obtained and incorporated into our analysis. For all sequenced cells, we counted the number of genes detected in each cell. Cells with fewer than 2,400 genes or 500,000 mapped reads were filtered out. In total, 80 oocytes and 71 GCs at five developmental stages passed the filter standards. To ensure the accuracy of estimated gene expression levels, only genes with FPKM > 1 in at least one cell were analyzed^58^. Expression levels of each gene were plus one then log2 transformed in the following analysis. Expression levels of each oocyte and GC were in Table S1.

### Principal Component Analysis (PCA)

The Seurat method was applied to analyze the single-cell data (77 unmatured oocytes, three matured MII oocytes and 71 GCs) to observe the whole clustering profile ^59^. Only highly variable genes (coefficient of variation>0.5) were used as inputs for PCA. And the marker genes in PCA plot were plotted by the FeaturePlot function in Seurat package. To complement the PCA clustering more accurately, we also clustered the oocytes and GCs separately using the FactoMineR package in R ^60^.

### Identification of DEGs and Gene Ontology Analysis

Multiple t-test was used to obtain the statistical significance of differentially expressed genes in each stage. Only the genes with significant p-values and false discovery rate (FDR) less than 0.05 with a fold change of log2 transformed FPKM larger than 1.5 were considered to be differentially expressed. Gene ontology analysis of differentially expressed genes was performed using DAVID ^61^.

### Identifying Expression Patterns of Maternal Effect Gene

Maternal effect genes play a critical role before zygote genome activation (ZGA). It’s reported that ZGA mainly happened at 4 cells to 8 cells stages ^30, 62^.To explore the expression patterns of maternal effect genes during folliculogenesis, we searched for genes carried from oocytes to early embryo and degraded before ZGA. Genes expressed (FPKM > 1) at MII and zygote stage but relatively low expressed (FPKM < 1) at 8 cells stage were taken as candidates. The candidate genes were clustered by expression correlation and cut into four clusters with cutree function in R.

### Identifying Secretory Protein Coding Signature Genes

To identify candidate biomarkers for predicting ovarian reserve, we focused on the oocyte- or GC-derived secretory proteins. The secretory protein encoding genes were downloaded from the Human Protein Atlas database (https://www.proteinatlas.org/). The GTEx v6 database (https://www.gtexportal.org) was utilized to obtain information on gene expression level in tissues of interest. We aimed to evaluate only protein secreting genes expressed in ovary and therefore we focused on genes that were previously identified only in gonads and could be present in brain, but not in any other tissues ^63^. We assumed that brain-blood (B-B) barrier prevents secretion of the proteins into systemic circulation and thus reduces confounding effect on peripheral levels of these proteins. This assumption was based on the previously observed high levels of AMH in brain, which supports the idea of restrictive function of B-B barrier64.

### Transcription Factor Network Construction

Transcription factors play a key role in regulating development. To find the driver factors and construct their regulatory network in each two consecutive stages, we used ARACNe to perform the regulatory network analysis as described previously ^17^. First, the 1,469 human transcription factors in AnimalTFDB ^25^ and gene expression matrix were taken as inputs for ARACNe-AP software ^25^. Then, viper package in R was used to visualize the transcription factors and their target genes in each consecutive stage ^65^. Regulators with p-values less than 0.01 were inferred as driver factors in each two consecutive stages.

### GSEA Analysis

To identify the significant enriched pathways in each two consecutive development stages, we used Gene Set Enrichment Analysis (GSEA, http://software.broadinstitute.org/gsea/index.jsp) to perform enrichment analysis, and the KEGG pathway was used ^66^. The gene sets that showed nominal p-value less than 0.05 were chosen as enriched.

### Analysis of Conservation Between Human and Mouse

Human and mouse homologous genes were downloaded from Vertebrate Homology Database (http://www.informatics.jax.org/homology.shtml). Housekeeping genes were obtained from Human Protein Atlas. Human oocytes expressed genes were overlapped with mouse oocytes expressed homologous genes to find genes expressed in both human and mouse oocytes, and the housekeeping genes were filtered out from the overlapped gene set. The retained genes were divided as homologous DEGs (genes in DEGs of human oocytes) and homologous non-DEGs. The genes involved in oocyte maturation defect were downloaded from Online Mendelian Inheritance in Man (OMIM) database (https://www.omim.org/) and incorporated in to analysis.

## Supplementary Figure Legends

**Figure S1.**
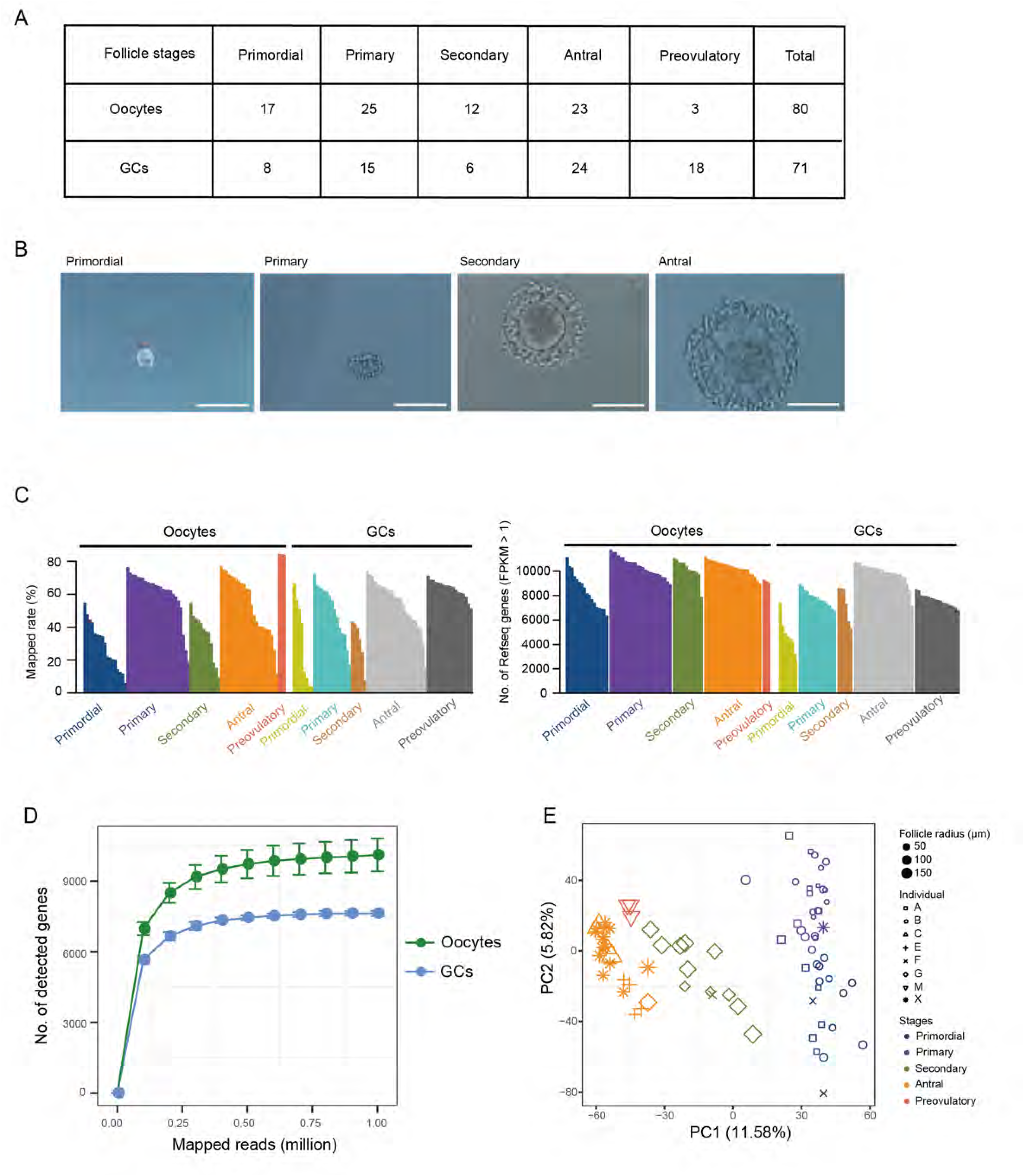
Sample Collection and Quality Control. Related to Figure 1 and 2. A) Table demonstrating number of samples per two follicle cell-type specific compartments at five follicle stages. B) Representation of the follicles in primordial, primary, secondary and antral stages.Scale bar, 100 µm. C) The number of detected genes (FPKM > 1) in each samples included in this study. D) Saturation analysis for scRNA-Seq data. X-axis is mapped reads for each down-sampling data, y-axis is number of genes detected. E) Principal component analysis (PCA) of the transcriptome of human oocytes that portrays sampling entities. Different colors and sizes of the points represent oocytes of different stages and sizes, respectively. Different shapes of the points represent different sampling entities (individuals).

**Figure S2.**
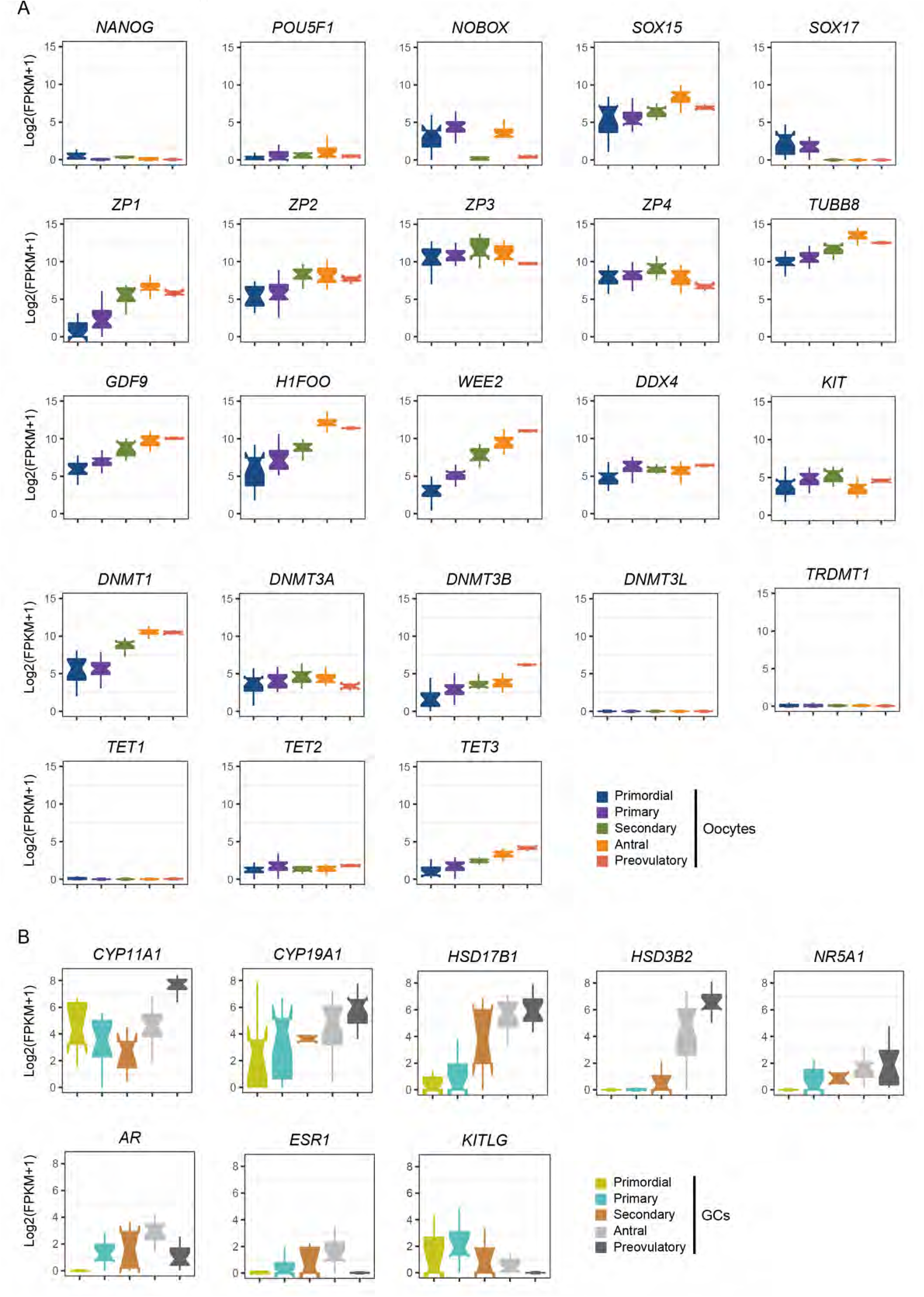
The Expression Patterns of Selected Marker Genes in Human Oocytes and GCs. Related to Figure 2 and 3. A) Boxplots of selected marker genes in oocytes at each follicle stage. B) Boxplots of selected hormone receptors and genes involved in steroidogenesis in GCs at each follicle stage.

**Figure S3.**
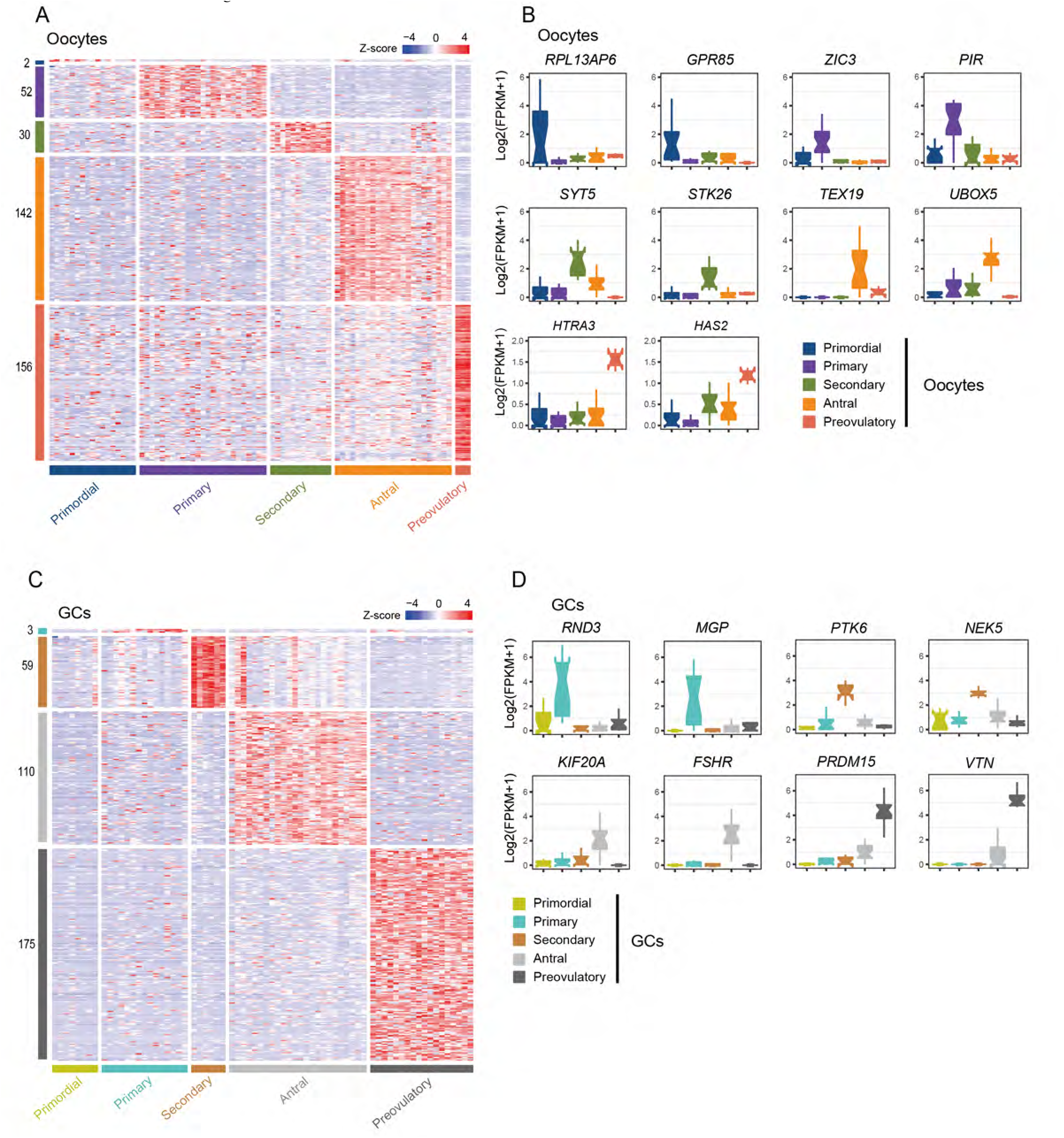
The Expression Patterns of Signature Genes in Oocytes and GCs at Each Stage. Related to Figure 2 and 3. A) Heatmap of signature genes in oocytes at five stages of folliculogenesis. The numbers of identified signature genes are indicated on the y-axis, and the stages of follicle development are presented along the x-axis. The color key from blue to red indicates the relative gene expression level from low to high, respectively. B) Boxplots of the relative expression levels (log2 [FPKM+1]) of selected signature genes in oocytes to exemplify the expression patterns in each stage. C) Heatmap of signature genes in GCs at different stages of folliculogenesis. D) Boxplots of the relative expression levels (log2 [FPKM+1]) of selected signature genes in GCs to exemplify the expression patterns in each stage.

**Figure S4.**
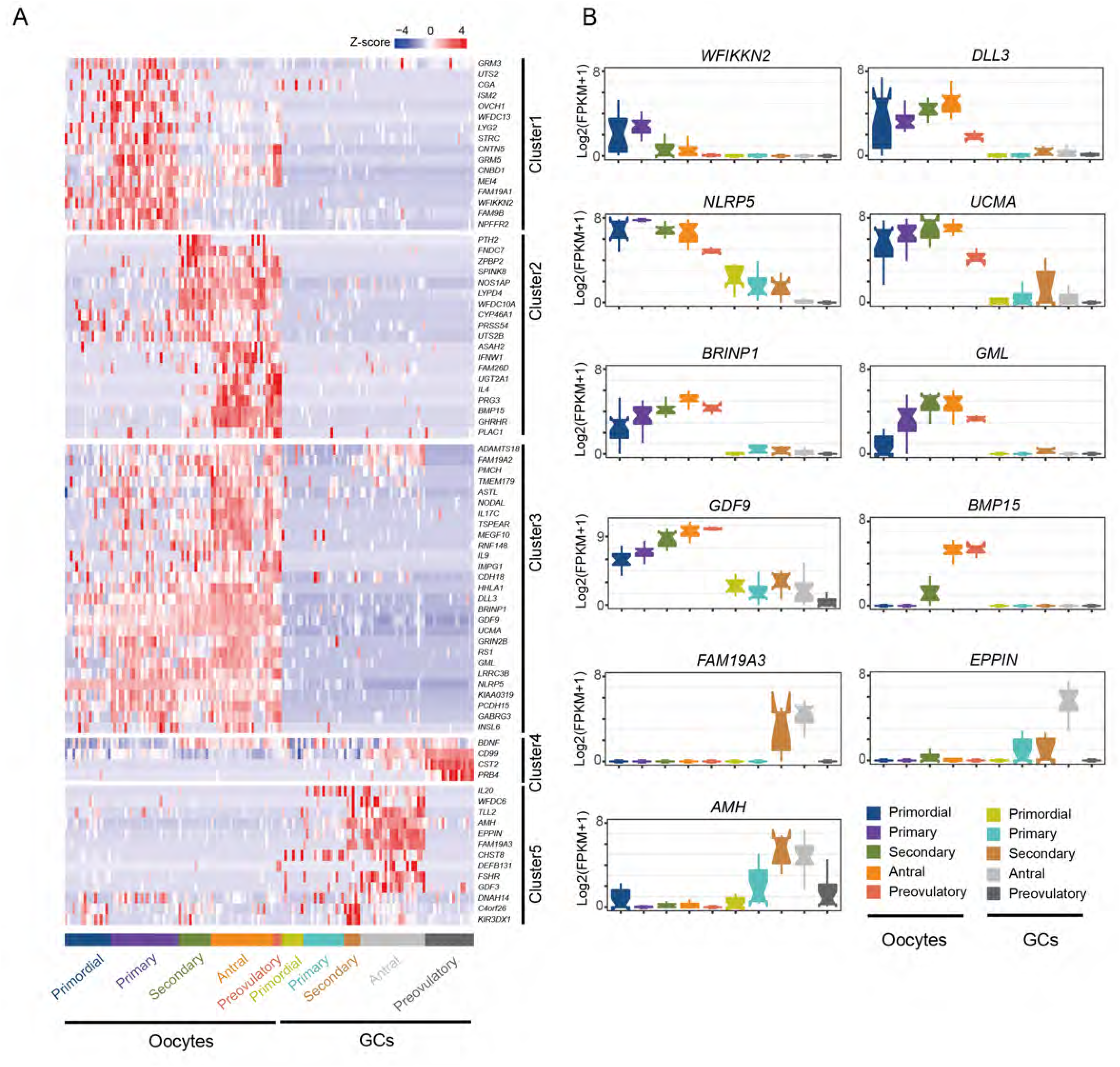
Secretory Protein Coding Signature Genes for Ovary Reserve Prediction. A) Heatmap of the secretory protein encoding genes expressed in oocytes (left) and GCs (right). The list of these genes is indicated on the y-axis according to the five clusters by similarity of expression patterns, and the stages of follicle development are presented along the x-axis. The color key from blue to red indicates the relative gene expression level from low to high, respectively. B) Boxplots of the relative expression levels (log2 [FPKM+1]) of selected secretory proteins encoding genes to exemplify the expression pattern in each cluster.

**Figure S5.**
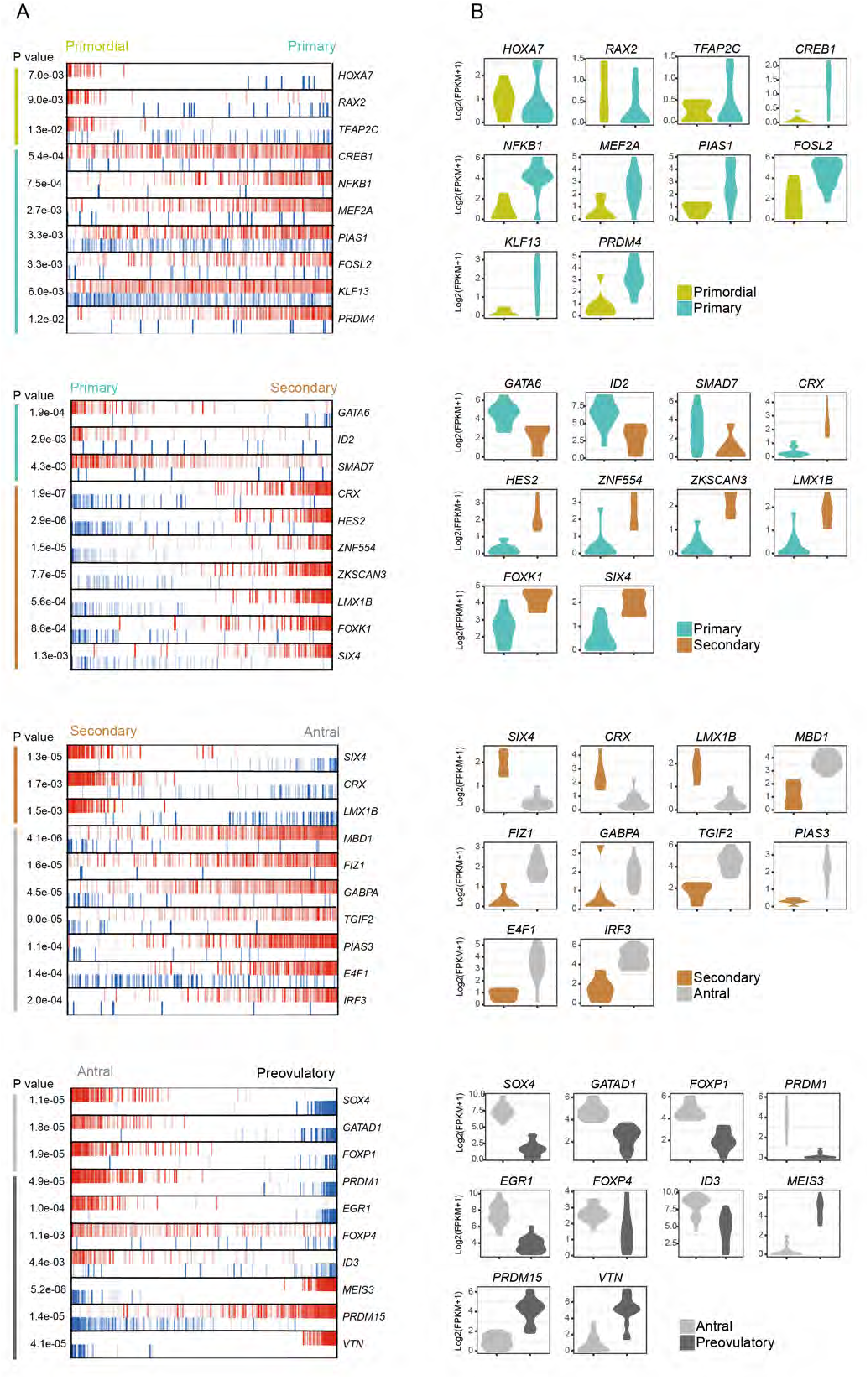
Inferred Key Transcriptional Factors in GCs at Each Stage-to-stage Transition of Folliculogenesis. Related to Figure 4. A) MARINa plots of targets for each candidate master regulator. Red vertical bar represents the activated targets, blue represents the repressed targets. On the x axis, genes were rank-sorted according to the significance of differential expression between the two developmental stages. The candidate master regulators were displayed on the right. The corresponding p-values of these master regulators indicate the significance of enrichment were displayed on the left. B) Violin plots show the relative expression levels (log2 [FPKM+1]) of each master regulator in two consecutive stages.

**Figure S6.**
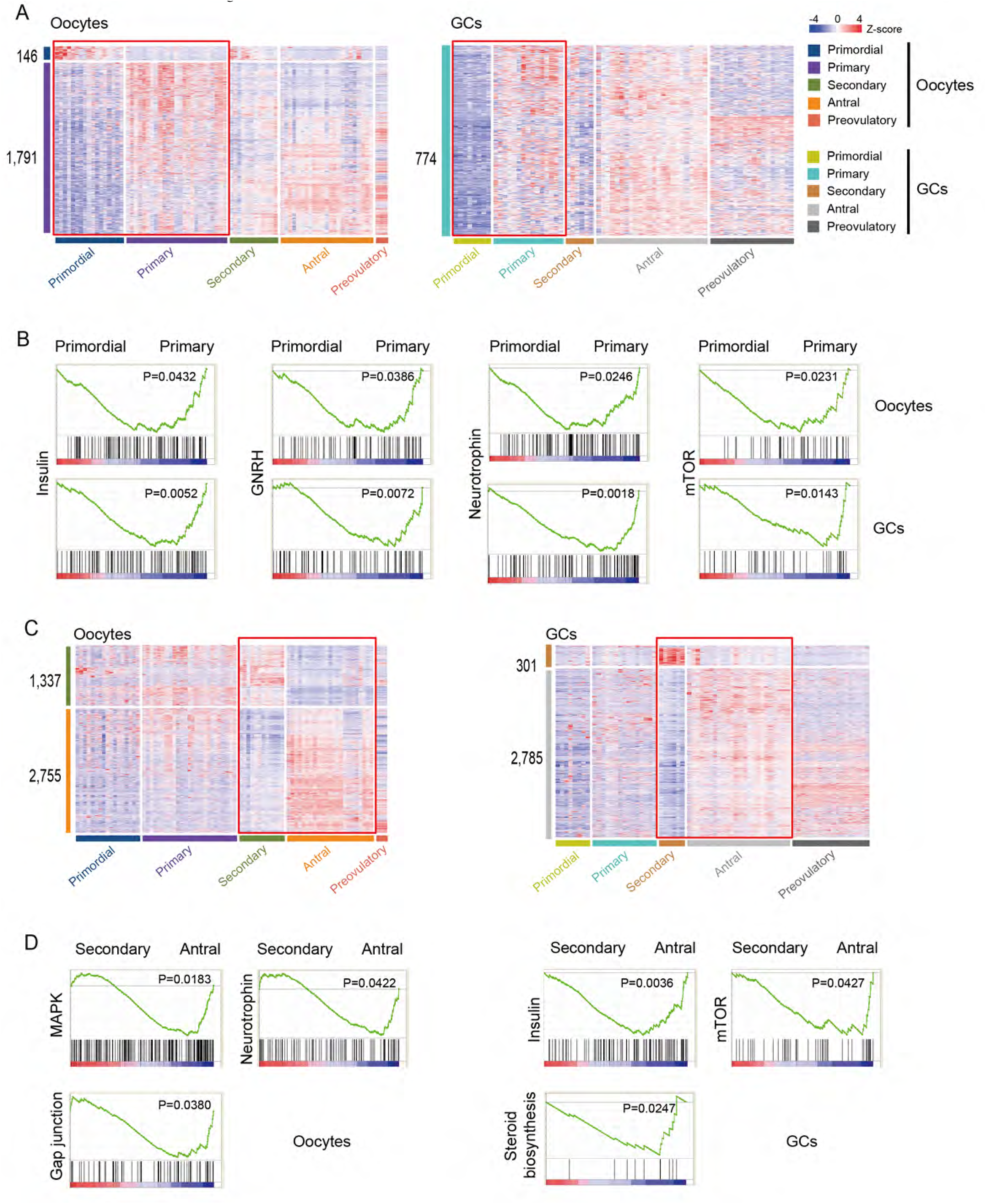
Signaling Pathways Enriched in Follicle Recruitment and Antral Formation by GSEA/KEGG Analysis. A) Heatmaps of DEGs between primordial and primary stage in oocytes and GCs. The numbers of identified DEGs are indicated on the y-axis, and the stages of follicle development are presented along the x-axis. The color key from blue to red indicates the relative gene expression level from low to high, respectively. B) GSEA enrichment plots of KEGG signaling pathways in oocytes and GCs between primordial and primary stage. C) Heatmaps of DEGs between secondary and antral stage in oocytes and GCs. D) GSEA enrichment plots of KEGG signaling pathways in oocytes and GCs between secondary and antral stage.

**Figure S7.**
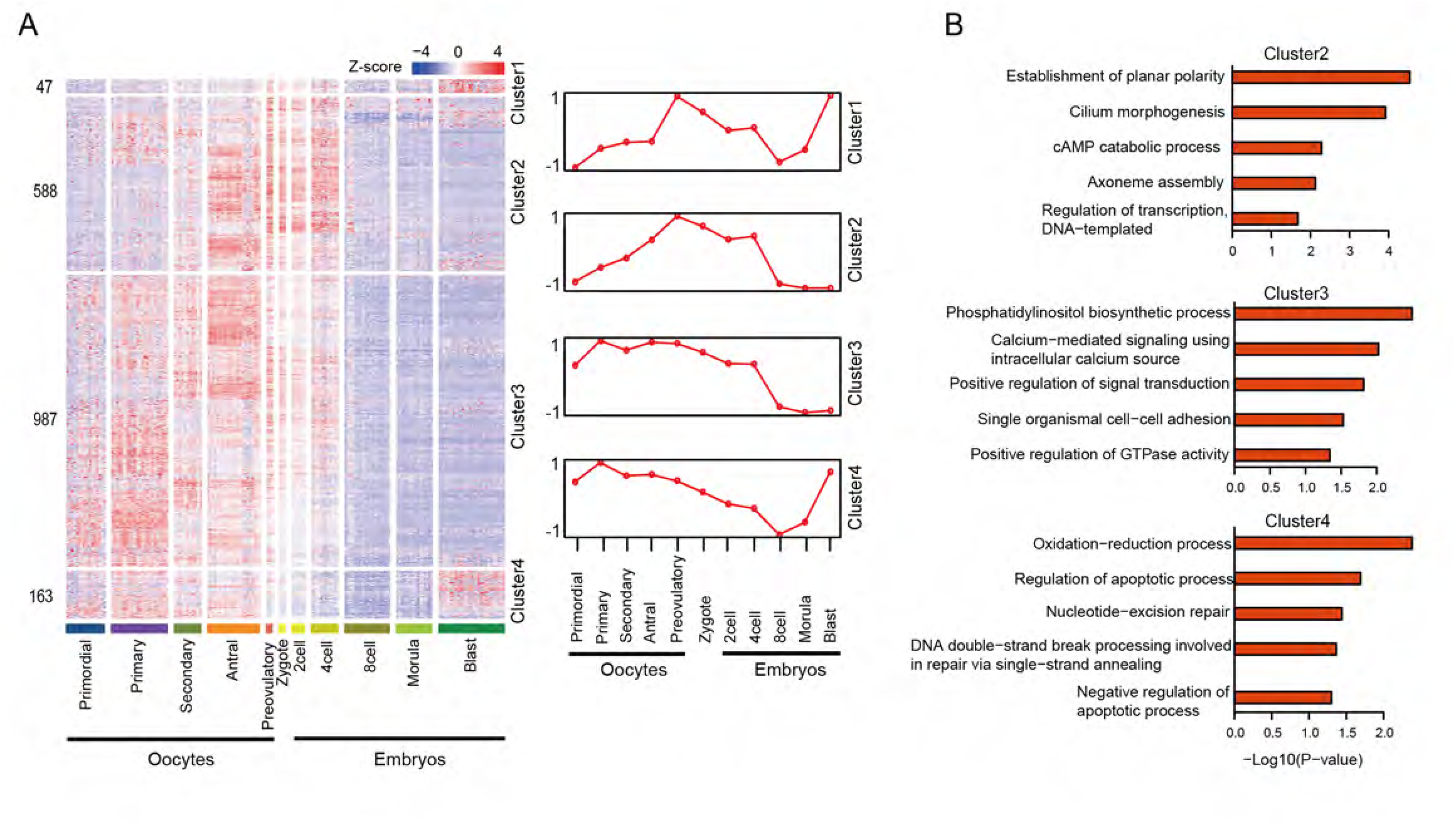
The Expression of Putative Maternal-Effect Genes in Folliculogenesis and Early Embryonic Development. A) Heatmap of the putative maternal-effect genes expressed in folliculogenesis and early embryonic development. The identified genes were clustered into four clusters by similarity of expression patterns. The stages of follicle development and early embryonic development are presented along the x-axis. Right panel show relative expression level of each cluster. The color key from blue to red indicates the relative gene expression level from low to high, respectively. B) Bar plots show the enriched GO terms of each cluster genes.

